# Specific targeting and clustering of Phosphatidylserine lipids by RSV M protein is critical for virus particle production

**DOI:** 10.1101/2023.03.13.532372

**Authors:** Jitendriya Swain, Maxime Bierre, Laura Veyrié, Charles-Adrien Richard, Jean-Francois Eleouet, Delphine Muriaux, Monika Bajorek

## Abstract

Human Respiratory Syncytial virus (RSV) is the leading cause of infantile bronchiolitis in the developed world and of childhood deaths in resource-poor settings. The elderly and the immunosuppressed are also affected. It is a major unmet target for vaccines and anti-viral drugs. RSV assembles and buds from the host cell plasma membrane by forming infections viral particles which are mostly filamentous. A key interaction during RSV assembly is the interaction of plasma membrane lipids with the Matrix (M) protein layer forming at assembly sites. Although the structure of RSV M protein dimer is known, it is unclear how the viral M proteins interact with certain plasma membrane lipids to promote viral assembly. Here, we demonstrate that M proteins cluster at the plasma membrane by selectively binding with phosphatidylserine (PS). Our *in vitro* studies suggest that M binds PS lipid as dimers and M mutant with a phosphomimetic substitution inhibits interaction with PS lipids. The presence of other negatively charged lipids like PI(4, 5)P2 does not affect RSV M ability to bind, while cholesterol negatively affects M interaction with lipids. Moreover, we show that the initial binding of the RSV M protein with lipids is independent of the cytoplasmic tail of fusion (F) glycoprotein (FCT). It is the first *in vitro* study of M interaction with plasma membrane lipids. M binding to plasma membrane may represent a viable therapeutic strategy for small molecules that will block viral spread.

## Introduction

Respiratory Syncytial virus (RSV) is a major public health issue. Human RSV is the most frequent cause of infantile bronchiolitis and pneumonia worldwide. In France, 460,000 infants are infected each year, and it is the first cause of hospitalization of young children. RSV hospitalization in elderly is comparable to influenza. The enormous burden of RSV makes it a major unmet target for a vaccine and anti-viral drug therapy (1). Recently, RSV vaccines for adults (Pfizer, GSK) and a mAb for infants were announced (AstraZeneca/Sanofi), but both miss important targets like infant vaccination and affordable therapies for low-income countries. The lack of knowledge of the RSV assembly and budding mechanism also presents a continuing challenge for large scale virus-like particle (VLPs) production for vaccine purposes. Therefore, understanding RSV assembly mechanism will open a new platform for both therapeutic strategy and vaccine development.

RSV belongs to the *Pneumoviridae* family in the order *Mononegavirales (2)*. The viral genome, encapsidated by the nucleoprotein (N), forms a ribonucleoprotein (RNP) complex, which constitutes the template for the viral polymerase. It was shown that RSV replication and transcription take place in virus-induced cytoplasmic inclusions called inclusion bodies (IBs) (3). According to the common paradigm, RSV assembles on the plasma membrane, and infectious viral particles are mainly filamentous (4, 5). However, more recent data suggest that viral filaments are produced and loaded with genomic RNA prior to insertion into the plasma membrane. According to this model, vesicles with RSV glycoproteins recycle from the plasma membrane and merge with intracellular vesicles, called assembly granules, containing the RNPs (6). RSV virions then assemble and bud (7) forming infectious viral particles which are mainly filamentous (8). The minimal set of RSV proteins required for viral filament formation are the cytoplasmic tail of fusion (F) glycoprotein (FCT), the phosphoprotein (P), and the matrix (M) protein (9, 10). M, a key structural protein, directs assembly, forming a protein lattice at specific assembly sites underlaying the plasma membrane. These M lattice supposedly bridges viral glycoproteins via the FCT and the internal RNP complex, the latter via binding to oligomeric N and/or P associated to the viral genome (10, 11). FCT was shown to be essential for viral filament formation, specifically the three last amino acids (Phe22-Ser23-Asn24) of the tail (12). A sole mutation Phe22Ala resulted in a complete loss of filament formation, possible due to loss of interaction with M or P (13). Recently, M and P alone were shown to form a sufficient platform for Virus-Like Particles (VLPs) budding, producing released particles that were highly variable in shape and size (14), suggesting that F is not required for the budding itself.

M is a main driver of RSV filament formation and budding (9, 15, 16). Previous structural data from our group showed that M forms dimers that are critical for viral filament assembly and VLP production (17). RSV M unstable dimers result in defects in higher-order oligomerization and as a consequence lack of filament formation and budding (17, 18). Additionally, M carrying phosphomimetic residue substitution that affects higher-order oligomer assembly leads to un-infectious virus production (5). Furthermore, recent cryoelectron tomography (cryo-ET) data has shown a helical lattice of M organized as dimers beneath the viral membrane and proposed that correctly ordered M layer may have implication on the conformation of the F protein, the principal target for vaccine and monoclonal antibodies (19).

Each monomer of M protein comprises two compact β-rich domains connected by an unstructured linker region. An extensive contiguous area of positive surface charge covering 600 Å^2^ and spanning both domains was suggested to drive the interaction with negatively-charged membrane surface. This positive region is complemented by regions of high hydrophobicity and a striking planar arrangement of tyrosine residues encircling the C-terminal domain, which make it suitable to target different phospholipids of plasma membrane surface (20).

Regardless of the cellular location, the interaction of M with specific lipids during assembly is not fully understood. Our previous works on enveloped RNA viruses – lipid interactions during virus assembly at the cell plasma membrane (21–25) reveal that retroviruses or influenza viruses always hijack specific phospholipids creating a platform for assembly, independently of envelope glycoproteins, but strongly dependent on Matrix residues (26). These interactions being mandatory for generating an enveloped viral particle. Similarly, the sorting of RSV M protein into lipid rafts has been found to be dependent on the presence of cell surface glycoproteins. However, in their absence, M protein is still present on the plasma membrane but not concentrated in lipid rafts (27).

*In vitro* studies have shown that RSV M protein interacts with lipid monolayers with neutral lipid compositions (DOPC/DPPC/cholesterol and DOPC/SM/cholesterol), and this interaction does not appear to be affected by the hydrophobic effect or the presence of cholesterol (28). More recent research has revealed that a specific set of lipids is required for M protein lattice formation in some paramyxoviruses, such as Nipah and measles (29). These viruses require phosphatidylserine (PS) and phosphatidylinositol-4,5-bisphosphate (PI(4,5)P2) for M protein interaction with lipids and M protein oligomerization. Overall, the lipid interactions of M proteins from different enveloped viruses appear to be diverse, but several common themes have emerged. Many of these M proteins associate with lipid rafts and interact with phospholipids, such as PS and PI(4,5)P2 (25, 29–32). These interactions appear to play a crucial role in the spread of these viruses and may provide a target for antiviral therapies.

In this work, we demonstrate that the interaction between RSV M protein and the host cell membrane is specifically facilitated by the PS lipid. Our findings show that the binding of M to lipids is not dependent on the presence of highly negatively charged lipids like PI(4, 5)P2 and that hydrophobic cholesterol negatively impacts initial M binding. Our results indicate that M alone can form clusters with PS lipids, and the interaction between M protein and lipids is not influenced by the FCT protein. Our study, based on M mutant proteins *in vitro*, also suggests that M binds PS lipids as dimers and that a M mutant with a phosphomimetic substitution inhibits interaction with PS lipids. Furthermore, we confirm that although M, P, and FCT are all necessary for the formation of RSV-like filaments, M and P are the minimum components required to form VLPs. This is the first demonstration of M interaction with physiologically relevant membrane lipids *in vitro* and can be used to further investigate the budding mechanism of RSV.

## Results

### M specifically interacts with negatively charged PS in the presence of neutral and PS lipid

Although it is known that RSV M binds plasma membrane lipids during assembly, it is unclear whether the interaction is unspecific towards negatively charged membrane surface or rather particular lipid head groups anchor M to the inner membrane leaflet. To study RSV M interaction with lipids, we used liposome sedimentation assays and large unilamellar vesicles (LUVs) to identify specific lipids critical for interaction (Fig. 1A). LUVs with Egg-Phosphatidylcholine (EPC) (100), or EPC:brain Phosphatidylserine (PS) at molar ratio of 70:30 were prepared as described in Experimental procedure section and incubated with M protein. M incubation without LUVs served as negative control. Samples were then centrifugated to separate the supernatant (S) fraction containing the unbound M and pellet (P) fraction containing the LUV-bound M. Both fractions were migrated on SDS gels and stained with Coomassie blue for visualization and quantification. As shown in Fig. 1B, about 20% of M was found in P fraction in the absence of LUVs. This agrees with previous results showing RSV M oligomerization and precipitation over time (17). This may also be a reason for different percentage of RSV M negative control found in P fraction with different set of experiments. In presence of PC LUVs similar percentage of M was found in the pellet indicating that no significant binding was observed. In contrast, when incubating with PC:PS LUVs, up to 50 %, which is 2.5 times more as compare to negative control, of M was found in the P fraction reflecting the PC:PS LUV-bound M (Fig. 1B). Our results show that RSV M specifically interacts with negatively charged PS in presence of neutral lipids like PC. Next, we investigated whether M could interact with lipids as dimer. We used an RSV M mutant, Y229A, which was previously shown to form dimers but be deficient in the formation of higher-order M oligomers and filaments (17). We incubated both M WT and M Y229A with PC:PS LUVs and then separated the S and P fractions by centrifugation. The samples were then analyzed on SDS gels and stained with Coomassie blue for visualization and quantification (Fig. 1C). In the absence of liposomes, negative control, approximately 11.3% and 10.1% of M WT and M Y229A were found in the P fraction, respectively. When incubated with PC:PS LUVs, an average of 31% and 26.35% of M WT and M Y229A were found in the P fraction, respectively, reflecting 2.7 times and 2.6 times more binding of M WT and M Y229A to lipids compared to the negative control. These results demonstrate that both RSV M proteins interact similarly with PC:PS liposomes, indicating that RSV M can interact with lipids as dimers and that M oligomerization is not necessary.

**Figure 1.**
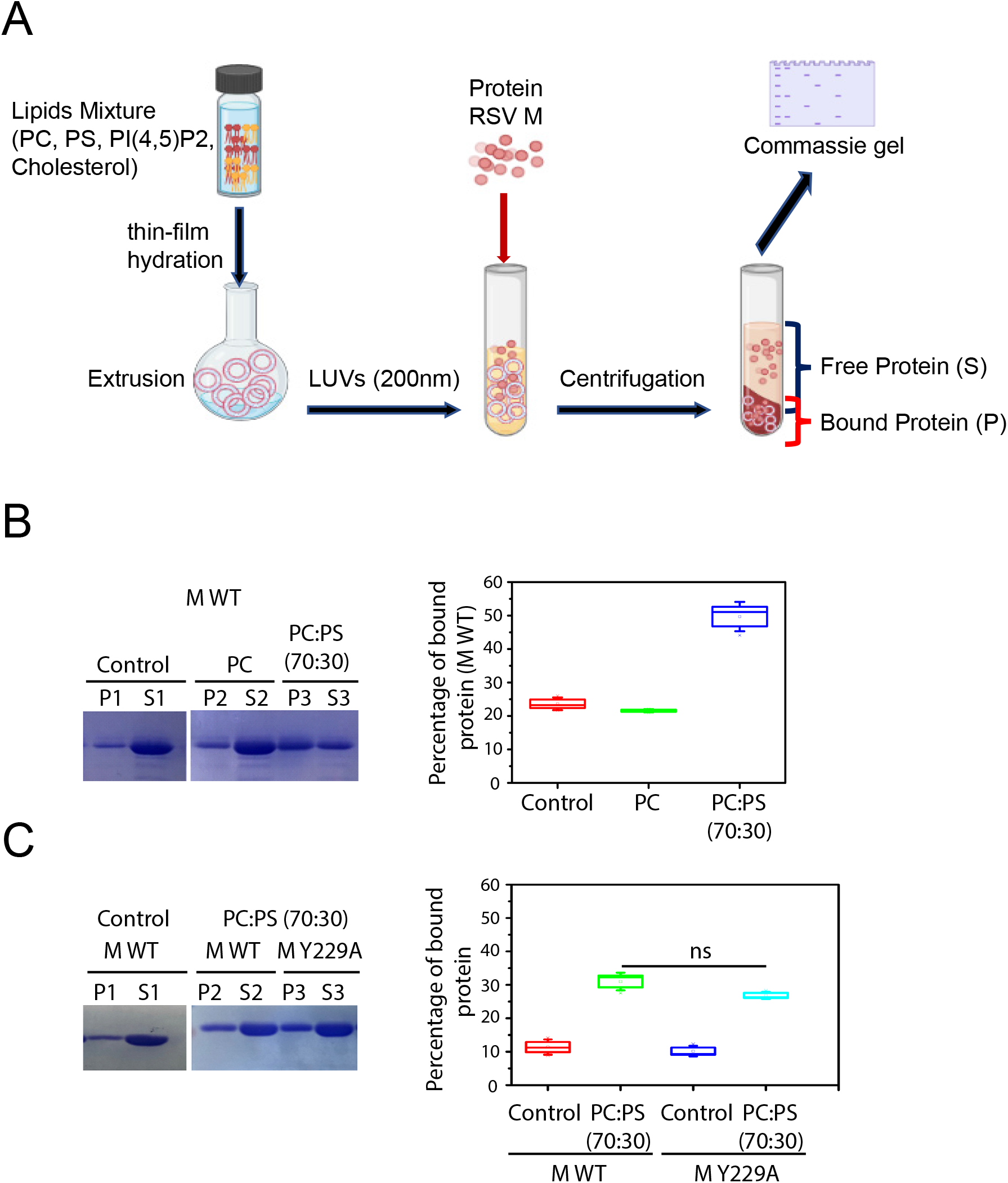
M specifically interacts with negatively charged PS. **A)** Schematic representation of co-sedimentation assay used. **B)** SDS PAGE obtained after co-sedimentation assay and staining with Coomassie blue showing RSV M protein negative control (P1,S1) (without any lipid), M with PC (P2,S2) and M with PC:PS(70:30) (P3,S3) lipid (pellet (P) and Supernatant (S)). The percentage of bound RSV M protein to lipids was quantified and is shown as a graph on the right. **C)** SDS PAGE obtained after cosedimentation assay and staining with Coomassie blue showing RSV M and M Y229A proteins negative control (P1,S1)(without any lipid), M with PC:PS (70:30) (P2,S2) and M Y229A with PC:PS (70:30) (P3,S3) lipid (pellet (P) and Supernatant (S)). The percentage of bound RSV M proteins to lipids was quantified and is shown as a graph on the right. Statistically significant analysis was evaluated using one□way ANOVA and t-parametric tests. *p* < 0.05 is significant. Full Coomassie gels for B and C are shown in Supp. Fig. 2.

### PI(4, 5)2 is not required and cholesterol negatively affects M binding to PS lipids

Recently, it was reported that Paramyxovirus Nipah and Measles M proteins interact with negatively charged PS lipids, and that PI(4,5)P2 significantly enhances this interaction (29). To examine the effect of PI(4,5)P2 on RSV M protein’s interaction with PS lipids, we conducted liposome sedimentation assays using PC:PS LUVs with and without PI(4,5)P2 (Fig. 2A). The results show that when incubated with PC:PS LUVs, M percentage increased in the P fraction (mean 29.81 %), reflecting LUV-bound M, compared to the negative control (mean 11.3 %). Similarly, when incubated with PC:PS:PI(4,5)P2 LUVs, the percentage of M in the P fraction (mean 28.14 %) increased 2.5 times compared to the negative control (Fig. 2A). This suggests that the presence of PI(4,5)P2 does not increase RSV M’s binding ability to lipid membranes.

**Figure 2.**
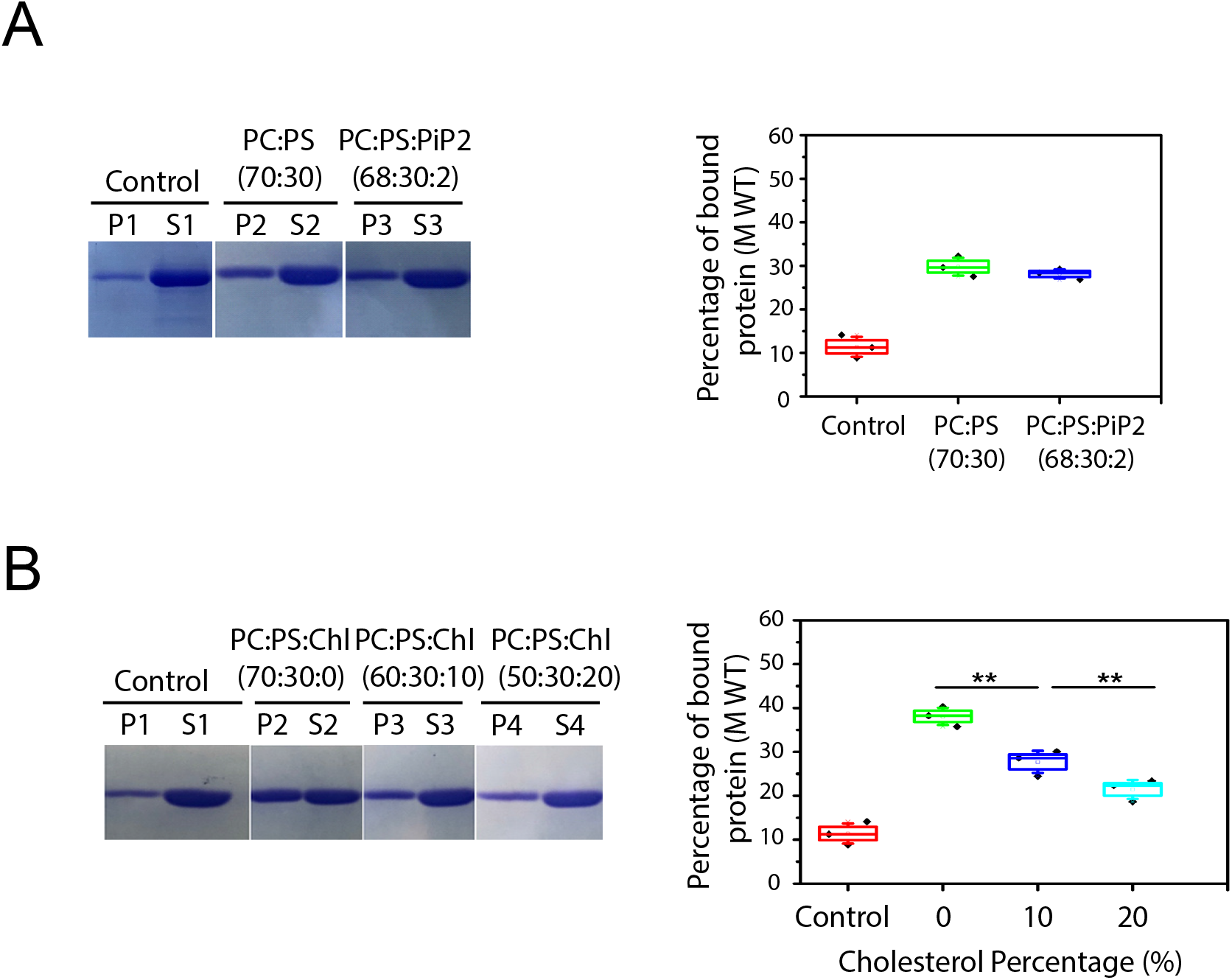
M protein interaction with PS in the presence of PI(4, 5)2 and cholesterol. **A)** SDS PAGE obtained after co-sedimentation assay and stained with Coomassie blue showing RSV M protein negative control (P1,S1) (without any lipid), PC:PS (70:30) (P2,S2) and PC:PS:PI(4,5)P2 (68:30:2) (P3,S3) lipid (pellet (P) and Supernatant (S)) and plot showing percentage of bound RSV M protein to lipids. **B)** SDS PAGE obtained after cosedimentation assay and stained with Coomassie blue showing RSV M proteins binding with different percentage of cholesterol, negative control (P1,S1) (without any lipid), M with PC:PS:Chl (70:30:0) (P2,S2), PC:PS:Chl (60:30:10) (P3, S3), and PC:PS:Chl (50:30:20) (P4,S4) lipid (pellet (P) and Supernatant (S)) and plot showing percentage of bound RSV M protein to lipids with or without cholesterol. Statistically significant analysis was evaluated using one□way ANOVA and t-parametric tests. *p* < 0.05 is significant. Full Coomassie gel for B is shown in Supp. Fig. 2.

The role of lipid rafts and Caveolae hydrophobicity is well established in RSV assembly and budding (27, 33, 34) but the specific role of cholesterol for M binding to lipids has not been investigated. Further, to study the effect of cholesterol on RSV M binding to lipids, we conducted sedimentation assays using PC:PS LUVs with and without different percentages of cholesterol (Fig. 2B). When M was incubated with PC:PS LUVs, 38.10% of M was found in the P fraction, which was 3.34 times more as compared to the negative control without LUVs (11.43%). However, incubation with PC:PS:Chl (60:30:10) and PC:PS:Chl (50:30:20) LUVs resulted in 27.7% and 21.4% of M in the pellet, respectively. These results show that cholesterol significantly reduces the binding ability of RSV M to PS lipids *in vitro*.

### The FCT protein is not required for RSV M protein-PS lipid binding on LUVs

The RSV FCT protein has been shown to be crucial for the production of infectious virus (13, 35). A lack of virus budding in FCT mutants has been suggested to be due to a loss of interaction with cellular or other RSV proteins, M or P (13). The RSV F protein is known to be a trimer (36), and the GCN4 protein (37) enables the trimerization of FCT, mimicking its natural state. We thus purified the trimeric FCT by using a His-GCN4-FCT construct. The purified construct migrates on SDS-PAGE according to its monomeric size of 8kD (Supp. Fig. 1). However, the sizing column profile clearly shows that His-GCN4-FCT protein forms a trimer migrating similarly to 30kD marker protein (Fig. 3A). To study the direct interaction between RSV M and FCT, different concentrations of DGS NTA-Ni lipid were added to PC:PS lipid mixtures to bind the His-GCN4-FCT protein to LUVs, showing an increase in binding (up to 80%) with an increase in the concentration of DGS Ni-lipid (Fig. 3B and 3C). The results from a sedimentation assay show that the FCT concentration significantly reduces the binding ability of M protein to PS lipids in presence of increasing concentration of FCT, decreasing from a mean of 38% to about 17% (Fig. 3B and 3D). This suggests that RSV M needs to directly interact with PS lipids, and its recruitment to lipids is not facilitated through FCT interaction.

**Figure 3.**
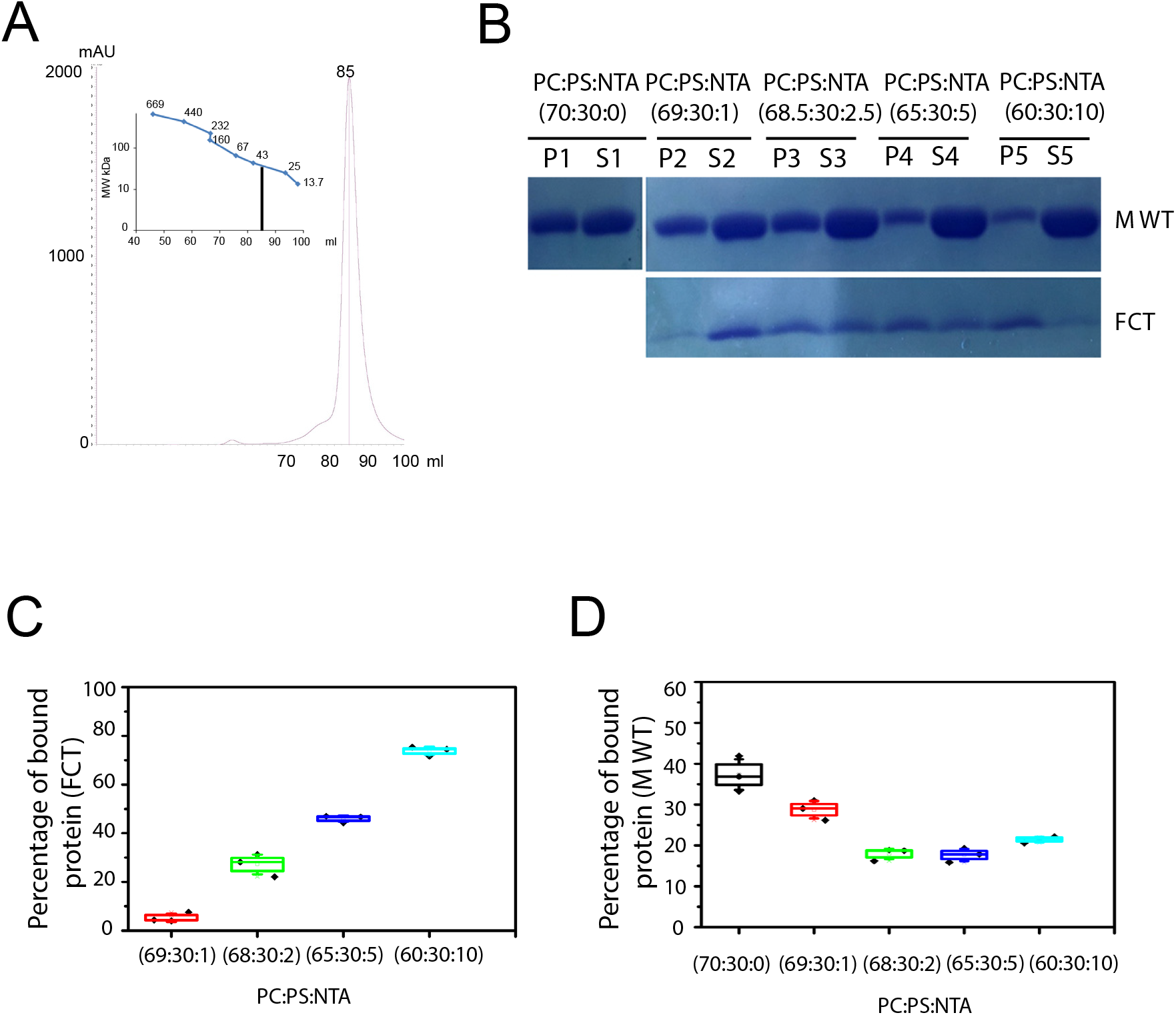
The FCT protein is not required for RSV M protein-PS lipid binding on LUV. **A)** Analytical size exclusion chromatography of His6-GCN4-FCT. Molecular mass of the FCT protein was estimated by comparing the gel-phase distribution of the FCT peak with the values obtained for known calibration protein standards (GE Healthcare). **B)** SDS PAGE obtained after co-sedimentation assay and stained with Coomassie blue showing RSV M and His6-GCN4-FCT protein binding PC:PS:NTA (70:30:0) (P1,S1), PC:PS:NTA (69:30:1) (P2,S2), PC:PS:NTA (68.5:30:2.5) (P3,S3), PC:PS:NTA (65:30:5) (P4, S4), and PC:PS:NTA (60:30:10) (P5, S5) lipid (pellet (P) and Supernatant (S)). Full Coomassie gel is shown in Supp. Fig. 2. **C)** A graph showing percentage of bound His6-GCN4-FCT protein incorporated to lipids **D)** A graph showing percentage of bound RSV M protein bound to lipids. Statistically significant analysis was evaluated using one□way ANOVA and t-parametric tests. *p* < 0.05 is significant

### RSV M protein induces clustering of PS lipid on model membranes *in vitro*

Our findings above demonstrate that RSV M binds to PS lipids with strong specificity and does not require the presence of other lipids and cholesterol (Fig. 1–3). Here, for the first time, we examined the clustering ability of M towards PS lipids on model membranes *in vitro*. To do this, we used a supported lipid bilayer (SLBs) containing 70% PC and 30% PS lipids, with a fluorescent PS lipid (Top Fluor-PS). Using time-lapse confocal imaging, we observed that RSV M quickly induced PS clusters on the SLBs (Supp. Movie 1). Fig. 4A shows the clustering of PS lipids with and without M protein: there were no or very few PS clusters in the absence of M, negative control. In contrast, adding 0.5 μM or 1 μM M induced PS cluster formation. We also measured PS cluster sizes and found that they increased in a concentration-dependent manner (Fig. 4B), from 0.16 μm^2^ to 2.86 μm^2^ with 0.5 μM and 1 μM of M respectively (after 30 min of incubation). Overall, our results show that RSV M is able to cluster PS on model membrane, probably a signature of RSV M oligomerization and assembly on model membranes through its interaction with PS.

**Figure 4.**
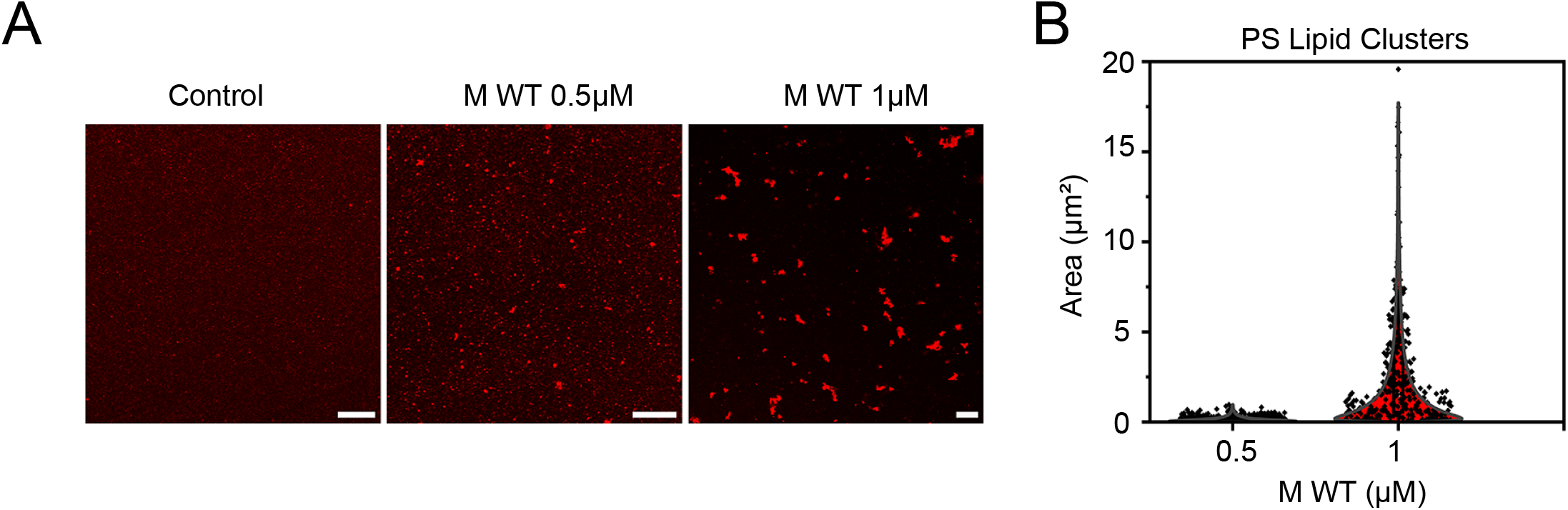
RSV M protein induces clustering of PS lipid. **A)** Time-lapse confocal imaging using SLBs containing 70% PC and 30% PS lipids, with a fluorescent PS lipid (Top Fluor-PS) with and without RSV M protein (0.5 μM and 1 μM). **B)** A graph showing PS clusters size in presence of RSV M protein. Statistically significant analysis was evaluated using one□way ANOVA and t-parametric tests. *p* < 0.05 is significant

### RSV M with T205D phosphomimetic substitution prevents interaction and clustering of PS lipids and generate abnormal virus-like filaments and VLPs

To assess whether M with a phosphomimetic substitution binds the same lipids like unphosphorylated M protein, we used a previously published M mutant, M T205D, which induces M higher-order oligomerization defects (5). We investigated whether M T205D interacts with PS lipids as we identified for WT M (Fig. 1–4). Again, we used the sedimentation assay and LUVs with PC:PS (7:3). When M WT was incubated with PC:PS LUVs, M again was found enriched in the P fraction (2.5 times more as compared to the negative control), indicating interaction with PS lipids. In contrast, when M T205D was incubated with the same PC:PS LUVs, there was no significant difference between the negative control protein found in the P faction and M T205D alone, suggesting no interaction (Fig. 5A). To further confirm this result, we analyzed the effect of M T205D on PS clustering using the SLBs PC:PS (70:30) system with the fluorescent PS lipid probe (Fig. 5B). There were no or very few PS clusters in the absence of M protein, negative control. No clusters formed when adding 1 μM M T205D protein. In contrast, adding 1 μM M WT induced PS cluster formation. Our results show that the M T205D mutant with phosphomimetic substitution is not able to induce clustering of the PS lipids *in vitro* on model membranes as compared to WT RSV M.

**Figure 5.**
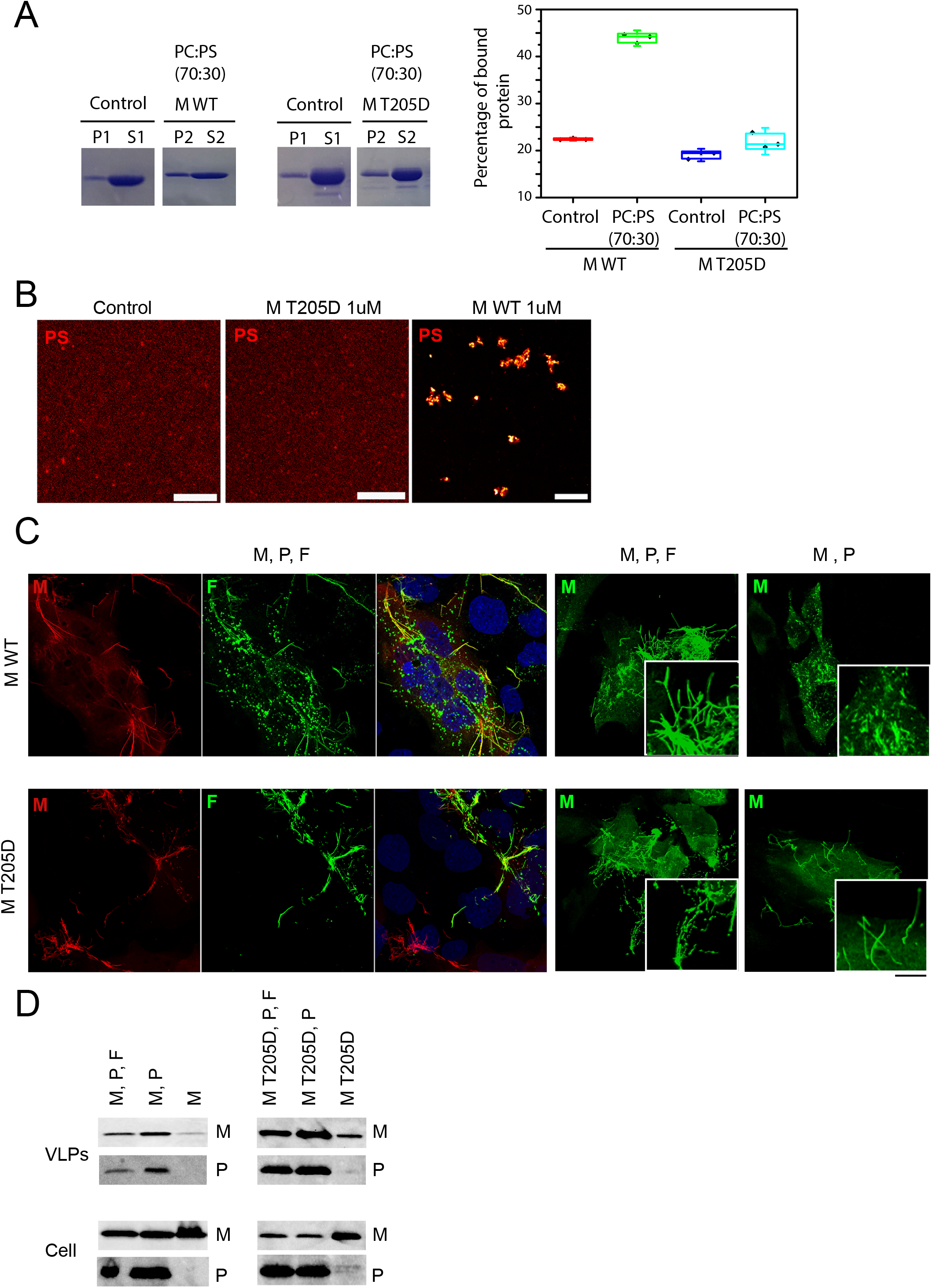
RSV M with phosphomimetic substitution on T205 prevents interaction and clustering of PS lipids and generate abnormal virus-like filaments and VLPs. **A)** SDS PAGE obtained after co-sedimentation assay and stained with Coomassie blue showing M protein negative control (P1,S1) (without any lipid), M with PC:PS (70:30) (P2,S2), M T205D negative control (P1,S1)(without any lipid), and M T205D with PC:PS (70:30) (P2,S2) lipid. A graph showing percentage of bound RSV M and M T205D protein to lipids is shown on the right. Full Coomassie gel is shown in Supp. Fig. 2. **B)** Time-lapse confocal imaging using SLBs containing 70% PC and 30% PS lipids, with a fluorescent PS lipid (Top Fluor-PS) without and with M or M T205D proteins (1 μM). **C)** BEAS-2B cells were co-transfected with pcDNA3.1 plasmids expressing RSV M WT or T205D mutant, P, and F or M and P alone. Cells were fixed, permeabilized at 24 h post transfection, immunostained with anti-M and anti-F or anti-M only primary antibodies followed by Alexa Fluor secondary antibodies, and were analysed by confocal microscopy. Scale bar represent 10 μm. Filaments are zoomed-in following anti-M staining. **D.** HEp-2 cells were cotransfected with pcDNA3.1 plasmids expressing RSV M WT (left panel) or T205D (right panel), P and F, or a different combination of these. At 48 h post transfection, cell lysates (bottom) were generated and VLPs (top) were isolated from the supernatant. VLPs and cell lysates were then subjected to Western analysis using anti-P and anti-M polyclonal antibody.

Recombinant RSV with M T205D mutation cannot be recovered (5). However, RSV filaments and VLPs can be generated independently of RSV infection by transfecting cells with plasmids encoding the M, P, and F proteins for virus-like filaments (10, 12) and M and P for VLPs (14) formation. We used this assay to compare the M WT and T205D mutant for their requirement of other viral proteins for filament and VLPs formation. BEAS-2B cells were transfected with M WT or T205D, P, with or without F, and the formation of RSV virus-like filaments was assessed using confocal imaging after co-staining with polyclonal anti-M and monoclonal anti-F antibodies (Fig. 5C, left panel) or staining with monoclonal anti-M antibody (Fig. 5C, right panel). Transfection of M WT, P and F (upper panel) resulted in virus-like filament formation as seen with anti-M staining and anti-F staining, as reported previously (10, 16), and the two proteins co-localized along the virus-like filaments (merged image). We then compared transfection with and without F followed by staining with another anti-M antibody. In contrast to transfection with M, P and F, transfection with M and P alone resulted in short protrusions only, without RSV-like filament formation, confirming that F is required for filament formation (12). Zoomed-in images are shown focusing on the filaments. In contrast, M T205D, P and F (lower panel) also formed virus-like filaments, but not all filaments were stained with anti-F antibody, suggesting that M T205D mutant can form viruslike filaments without F. This was confirmed by transfecting cells with M T205D and P alone. Staining with anti-M revealed filaments, although they seemed more branched and disorganized, as previously reported (5). Importantly, in contrast to M WT, M T205D mutant also formed virus-like filaments without F.

Next, we performed a VLP budding assay (Fig. 5D). HEp-2 cells were transfected to express WT or T205D M alone or with P and F expressing plasmids. Cell lysates (soluble fraction) and the VLPs released into the cell supernatant were analyzed by Western blotting using anti-M and anti-P antibodies. Transfecting cells with M, P and F resulted in the release of VLPs for both WT and T205D M. In the presence of M and P alone, VLPs were also detected, confirming that the two proteins form the minimal platform for RSV VLP budding (14). In contrast to M WT, the expression of M T205D alone resulted in the production of an abnormal VLPs (Fig. 5D) containing just M. Our results here show that M T205D prevents interaction and clustering of PS lipids, and forms abnormal virus-like filaments and VLPs.

## Discussion

In this work, we found that PS lipids mediate the interaction between RSV M protein and host membrane (Fig. 1). It was previously proposed that an extensive contiguous area of positive surface charge on the M monomer covering 600 Å^2^ and spanning both N- and C-terminal domains drives the interaction with any negatively charged membrane surface (20). Our work here shows that the interactions between the RSV M protein and the lipid bilayer are indeed electrostatic in nature, but we also specifically identified the PS as the main drive for M binding to lipid at the plasma membrane. We also show that RSV M is able to cluster PS lipids on model membrane (Fig. 4), probably a signature of RSV M oligomerization and assembly through its interaction with PS.

In comparison to the RSV M protein, several other viruses also have matrix proteins that bind to negative lipids on plasma membranes and induce lipid clustering. For example, the human immunodeficiency virus (HIV) matrix protein has been shown to bind to lipids such as PS and, more specifically, PI(4,5)P2 and induce cluster formation (25). It has been demonstrated that also the matrix proteins of the Ebola, Marburg, Paramyxovirus Nipah, and Mesaels viruses more specifically bind to PI(4,5)P2 at plasma membranes and cause lipid clustering (29, 31, 32). Surprisingly, the presence of other negatively charged lipids like PI(4, 5)P2 does not affect RSV M ability to bind, and this ability is highly specific to PS lipids (Fig. 2A). In fact, the RSV M protein interacts similarly to the M1 matrix protein of influenza A virus, which only interacts with PS lipid and produces lipid clusters (23, 30).

We have also investigated whether the presence of cholesterol negatively affects initial M binding to lipids (Fig. 2B). The previous data is somehow controversial. In RSV infected cells, M was shown to be associated with lipid rafts (33, 38, 39). Moreover, M interacts with Caveolae proteins, Cav-1 and Cav-2 (40), which are the major components of lipid rafts together with cholesterol. However, the sorting of M into lipid rafts was shown to be dependent on the presence of cell surface glycoproteins. In the absence of glycoproteins, M was still found on the plasma membrane but not concentrated in lipid rafts (27). Our results, showing that increasing cholesterol concentration in LUVs prevents M interaction with lipids (Fig. 2B), may reflect the initial M binding to plasma membrane, which is independent of lipid rafts. Although RSV budding, when all the viral proteins assemble together, is most likely to occur at lipid rafts, the initial M binding may occur elsewhere on the plasma membrane, most probably at PS lipids like shown in Fig. 1–4.

Earlier studies using Cryo-EM of culture-grown RSV have determined the architecture of the virus. The presence of an intact M layer beneath the viral membrane was linked to the virion’s pre-fusion F form (8). Most recent cryo-electron tomography (cryo-ET) data has shown a helical lattice of M organized as dimers beneath the viral membrane, further confirming that M has implications on the conformation of the F protein (19). These M lattices were suggested to bridge viral glycoproteins via the FCT and the internal RNP complex, the latter via binding to oligomeric N and/or P associated with the viral genome (10, 11). Moreover, FCT was shown to be required for infectious virus production; however, viral filaments in FCT mutants still formed occasionally and contained M (12). In another study FCT was also shown to be essential for budding, specifically the three last amino acids (Phe22-Ser23-Asn24) of the tail (13). Overall, M interaction with FCT was often suggested but never demonstrated directly. Here, we show that the initial binding of the M protein with PS lipids is independent of the FCT protein (Fig. 3). This is also in agreement with previously published data (14) and our results in Fig. 5D, which show M and P being a minimal set for VLP production. M is found in the VLPs in the absence of F. The helical lattice of M organized as dimers beneath the viral membrane in close proximity to F tails seen in the virus suggests M interaction with FCTs but this most probably occurs later and is not required during budding. We cannot exclude that the GCN4-FCT construct, made to mimic the trimeric FCT, is not properly positioned regarding the distance from the lipid layer since the GCN4 domain results in distancing the FCT from the membrane. However, if the GCN4-FCT accurately mimics the trimeric FCT, the results strongly suggest that M must initially bind membranes independently of FCT.

Our results based on RSV M oligomerization mutant proteins *in vitro* suggest that M binds PS lipids as dimers (Fig. 1C). To our knowledge, the question of whether M binds plasma membranes as dimers or as oligomers was not addressed previously. The basic RSV M unit is a dimer (17) and M in the cytoplasm was shown to be dimeric (18). This makes sense, since M is also found in IBs which are membrane free viral structures (3, 41). We therefore conclude, based on our results and published work, that M binds to plasma membrane lipids as dimers and only forms higher-order oligomers during viral filament formation and VLPs budding. Similar results were shown also for other viral Matrix proteins, for example EBOV and MARV VP40 (42).

M was shown to be phosphorylated in infected cells as early as 6 h post infection (43). Our work here shows that M mutant, T205D, with phosphomimetic substitution prevents interaction and clustering of PS lipids (Fig. 5A and 5B). However, T205D forms virus-like filaments, but abnormal (Fig. 5C), and can bud on its own (Fig. 5D). This indicates it is able to bind to plasma membrane, but possibly due to wrong protein-lipid interactions. Recombinant M protein produced in bacteria, which was used for *in vitro* LUVs sedimentation assays, is unphosphorylated. Since M T205D has a phosphomimetic substitution and does not bind PS lipids, we can speculate that M binds PS lipids at the plasma membrane as an unphosphorylated protein. We hypothesize that constantly phosphorylated M on T205 binds unspecific phospholipids, this induces defective oligomerization that results in branched and shorter viral filaments that bud without other viral proteins. This could explain why RSV carrying the M T205D mutation could not be recovered (5).

In conclusion, the results presented above demonstrate the specific binding of the RSV M protein to PS lipids and its ability to induce lipid clustering, similar to other enveloped viruses, highlighting the importance of this interaction for the virus. Understanding the mechanism of this interaction can have important implications for the development of new therapeutic strategy that aim to block viral spread.

## Experimental procedures

### Cell culture

HEp-2 cells were maintained in Dulbecco modified Eagle medium (eurobio) supplemented with 10% fetal calf serum (FCS; eurobio), 1% L-glutamine, and 1 % penicillin streptomycin. The transformed human bronchial epithelial cell line (BEAS-2B) (ATCC CRL-9609) was maintained in RPMI 1640 medium (eurobio) supplemented with 10% FCS, 1% L-glutamine, and 1% penicillin-streptomycin. The cells were grown at 37°C in 5% CO_2_.

### Plasmids

pCDF plasmids encoding RSV M or GCN4-FCT proteins were used for the expression and purification of recombinant M and GCN4-FCT proteins, and pcDNA3.1 codon-optimised plasmids encoding the RSV M, P and F proteins (gift from M. Moore, Emory University) were used for the expression of viral proteins in cells. M Y229A and MT205D substitutions were generated using the Quick change directed mutagenesis kit (New England Biolabs), as recommended by the manufacturer.

### Bacteria expression and purification of recombinant proteins

For M expression (WT, Y229A and T205D mutant), *E. coli* Rosetta 2 bacteria transformed with the pCDF-M plasmid were grown from fresh starter cultures in Luria-Bertani (LB) broth for 5 h at 32°C, followed by induction with 0.4 mM isopropylthi-galactoside (IPTG) for 4 h at 25°C. Cells were lysed by sonication (4 times for 20 s each time) and lysozyme (1 mg/ml; Sigma) in 50 mM NaH_2_PO_4_-Na_2_HPO_4_, 300 mM NaCl, pH 7.4, plus protease inhibitors (Roche), RNase (12 g/ml, Sigma), and 0.25% CHAPS {3-[(3-cholamidopropyl)-dimethylammonio]-1-propanesulfonate}. Lysates were clarified by centrifugation (23,425 g, 30 min, 4°C), and the soluble His6-M protein was purified on a Nickel sepharose column (HiTrap™ 5 ml IMAC HP; GE Healthcare). The bound protein was washed extensively with loading buffer plus 25 mM imidazole and eluted with a 25 to 250 mM imidazole gradient. M was concentrated to 2 ml using Vivaspin20 columns (SartoriusStedimBiotec) and purified on a HiLoad 10/600 Superdex S200 column (GE Healthcare) in 50 mM NaH_2_PO_4_-Na_2_HPO_4_, 300 mM NaCl, pH 7.4. The M peak was concentrated to 3 mg/ml using Vivaspin4 columns. The His tag was digested during 14h at 4°C with histidine-tagged 3C proteases (Thermo Scientific) according to the manufacturer’s recommendations, incubated for 40 min at 4°C with nickel phosphate loaded Sepharose beads (GE Healthcare) to separate the His tag, and concentrated using 10,000 MWCO Vivaspin20 columns. The quality of protein samples was assessed by SDS-PAGE. Protein concentration was determined by measuring absorbance at 280 nm.

For expression of recombinant His-GCN4-FCT, *Escherichia coli* BL21 bacteria were transformed with pCDF-FCT plasmid, grown in LB broth for 5 h at 32°C and then induced with 0.5mM final IPTG for 4 h at 25°C. Bacteria were resuspended in lysis buffer (50mM NaH2PO4-Na2HPO4, 300mM NaCl, pH 7.2) supplemented with 1mg/ml lysozyme (Sigma) and 10mg/ml protease inhibitors (Roche) and then lysed by sonication (45 seconds amplitude 50). The lysates were clarified by centrifugation (30 min at 23,425 g, 4°C) and the soluble proteins with His6 tag were purified on 1ml of nickel phosphate loaded sepharose beads (GE Healthcare). Bound proteins were washed in lysis buffer with 25mM imidazole then with 400mM Imidazole, eluted with 800mM imidazole, and concentrated to 2ml using Vivaspin4 5,000MWCO columns (Sartorius Stedim Biotec). His-GCN4-FCT was further purified on a HiLoad 10/600 Superdex S200 column (GE Healthcare) in 50mM NaH_2_PO_4_-Na_2_HPO_4_, 300mM NaCl, pH 7.2. Proteins were concentrated to 2mL final using Vivaspin4 5,000 MWCO columns.

Purification profiles of M WT, M Y229A, M T205D and His6-GCN4-FCT protein are shown in Supplamental Fig. 1

### *In vitro* co-sedimentation assays with large unilamellar vesicles (LUVs)

Binding of RSV M WT and mutant protein to different lipids was determined by cosedimentation assays with LUVs. LUVs were made at desired ratio of a mixture of Egg-Phosphatidylcholine (EPC), brain Phosphatidylserine (PS), Phosphatidylinositol-4,5-biphosphate (PI(4,5)P2) and cholesterol as per experimental condition. All lipids were purchased from Avanti Polar Lipids. Lipid mixtures were solubilized in chloroform and evaporated with rotavapor. Lipids were then resuspended with Tris-NaCl buffer (150mM NaCl, 10mM Tris-HCl, pH 7.4) using freeze-thaw cycles, and extruded with filter to obtain LUVs of 200nm. A required amount of desired protein was incubated with LUVs (1 mg/ml) in a final volume of 100μL at room temperature for 30 min. Samples were then centrifuged at 220,000 g in a Beckman TLA 100 rotor at 4°C for 30 min. Each sample was then divided in supernatant (S = 90 μl), containing unbound protein, and pellet (P = 10 μl), containing LUV-bound protein. P was diluted in 80 μl of Tris-NaCl buffer (150mM NaCl, 10mM Tris-HCl, pH 7.4) buffer to maintain the equivalence between the S and P volumes. Then, 20 μl of S and P were analyzed by SDS-PAGE and protein were detected by staining with Coomassie Blue. For Ni-incorporated LUVs preparation, we have used 1,2-dioleoyl-sn-glycero-3-[(N-(5-amino-1-carboxypentyl) iminodiacetic acid) succinyl] (nickel salt) lipids. The protein intensities (Is, Ip) were quantified using the Image J software. The percentage of LUV-bound protein was calculated as: % protein LUV-bound = 100*I_P_/(I_P_+I_S_).

### Supported lipid bilayer (SLBs) preparation

Vesicle fusion method for SLBs was used, as previously shown (44). The cover slips were cleaned with piranha solution (3:1 Sulphuric acid (H2SO4): Hydrogen peroxide (H2O2). 0.5 mg/ ml of LUVs was deposited on the cleaned glass slide for SLBs preparation, kept in 55°C for 30 min and then washed with Tris-NaCl (150mM NaCl, 10mM Tris-HCl, pH 7.4) buffer to clean unfused vesicles. Zeiss LSM980 confocal microscope was used at a nominal magnification of 63X oil for SLBs imaging.

### Virus-like filament/particle formation

Over-night cultures of BEAS-2B cells seeded at 4 10^5^ cells/well in 6-well plates (on a 16-mm micro-cover glass for immunostaining) were transfected with pcDNA3.1 codon-optimized plasmids (0.4 μg each) carrying the RSV A2 WT or T205D M along with pcDNA3.1 codon-optimized plasmids carrying RSV A2 P and F using Lipofectamine 2000 (Invitrogen) according to the manufacturer’s recommendations. Cells were fixed 24 h post transfection, immunostained, and imaged as described below. For VLP formation, over-night cultures of HEp-2 cells seeded at 4 10^5^ cells/well in 6-well plates were transfected as described above. Released VLPs were harvested from the supernatant; the supernatant was clarified of cell debris by centrifugation (1,300 g, 10 min, 4°C) and pelleted through a 20% sucrose cushion (13,500 g, 90 min, 4°C). Cells were lysed in radio immune precipitation assay (RIPA) buffer. Cellular lysates and VLP pellets were dissolved in Laemmli buffer and subjected to Western analysis.

### SDS-PAGE and Western analysis

Protein samples were separated by electrophoresis on 12% polyacrylamide gels in Trisglycine buffer. All samples were boiled for 3 min prior to electrophoresis. Proteins were then transferred to a nitrocellulose membrane (RocheDiagnostics). The blots were blocked with 5% non fat milk in Tris-buffered saline (pH 7.4), followed by incubation with rabbit anti-P antiserum (1:5,000) (45), rabbit anti-M antiserum (1:1,000), and horseradish peroxidase (HRP)-conjugated donkey anti-rabbit (1:10,000) antibodies (P.A.R.I.S.). Western blots were developed using freshly prepared chemiluminescent substrate (100 mM TrisHCl, pH 8.8, 1.25 mM luminol, 0.2 mM p-coumaric acid, 0.05% H_2_O_2_) and exposed using BIO-RAD ChemiDoc™ Touch Imaging System.

### Generation of M antiserum

Polyclonal anti M serum was prepared by immunizing a rabbit three times at 2 week intervals using purified His-fusion proteins (100 mg) for each immunization. The first and second immunizations were administered subcutaneously in 1 ml Freund’s complete and Freund’s incomplete adjuvant (Difco), respectively. The third immunization was done intramuscularly in Freund’s incomplete adjuvant. Animals were bled 10 days after the third immunization.

### Immunostaining and imaging

Cells were fixed with 4% paraformaldehyde in PBS for 10 min, blocked with 3% BSA in 0.2% Triton X-100–PBS for 10 min, and immunostained with monoclonal anti-M (1:200; a gift from Mariethe Ehnlund, Karolinska Institute, Sweden), or double stained with polyclonal anti-M antiserum (1:1000) and monoclonal anti-F (1:500, BIO-RAD) antibodies, followed by species-specific secondary antibodies conjugated to Alexa Fluor 488 and Alexa Fluor 568 (1: 1,000; Invitrogen). Images were obtained using the White Light laser SP8 (Leica Microsystems, Wetzlar, Germany) or the Zeiss LSM700 confocal microscope at a nominal magnification of 63X oil. Images were acquired using the Leica Application Suite X (LAS X) software.

This article contains supporting information.

## Acknowledgment

These studies were supported by University of Montpellier to JS, by CNRS/UM to DM and by INRAE to JFE and MB.

