## Supplemental Figures for "Specific targeting and clustering of Phosphatidylserine lipids by RSV M protein is critical for virus particle production"

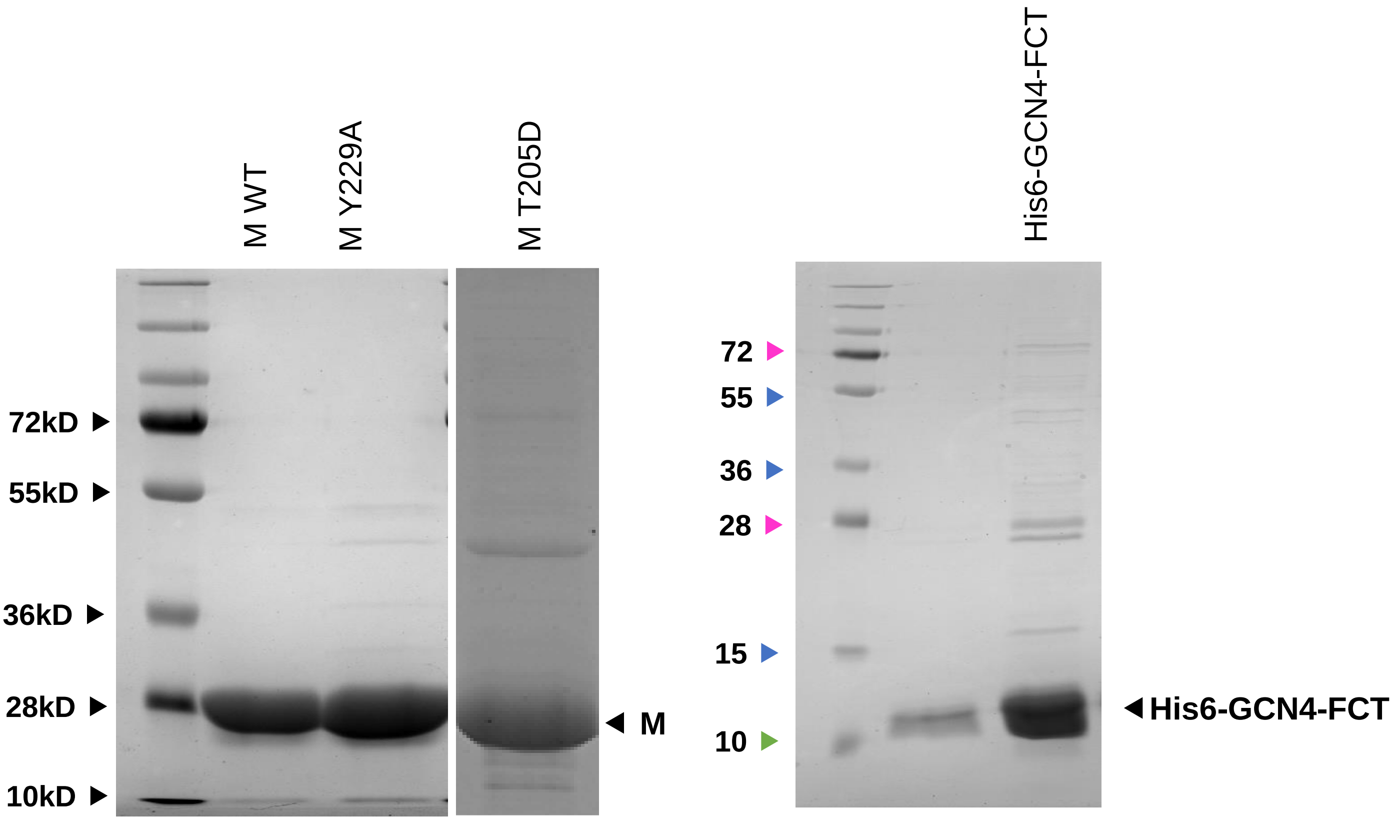


**Supplemental Figure 1**: Purification profiles for M and FCT proteins purified on Nickel sepharose and HiLoad 10/600 Superdex S200 column as described in Experimental procedures .

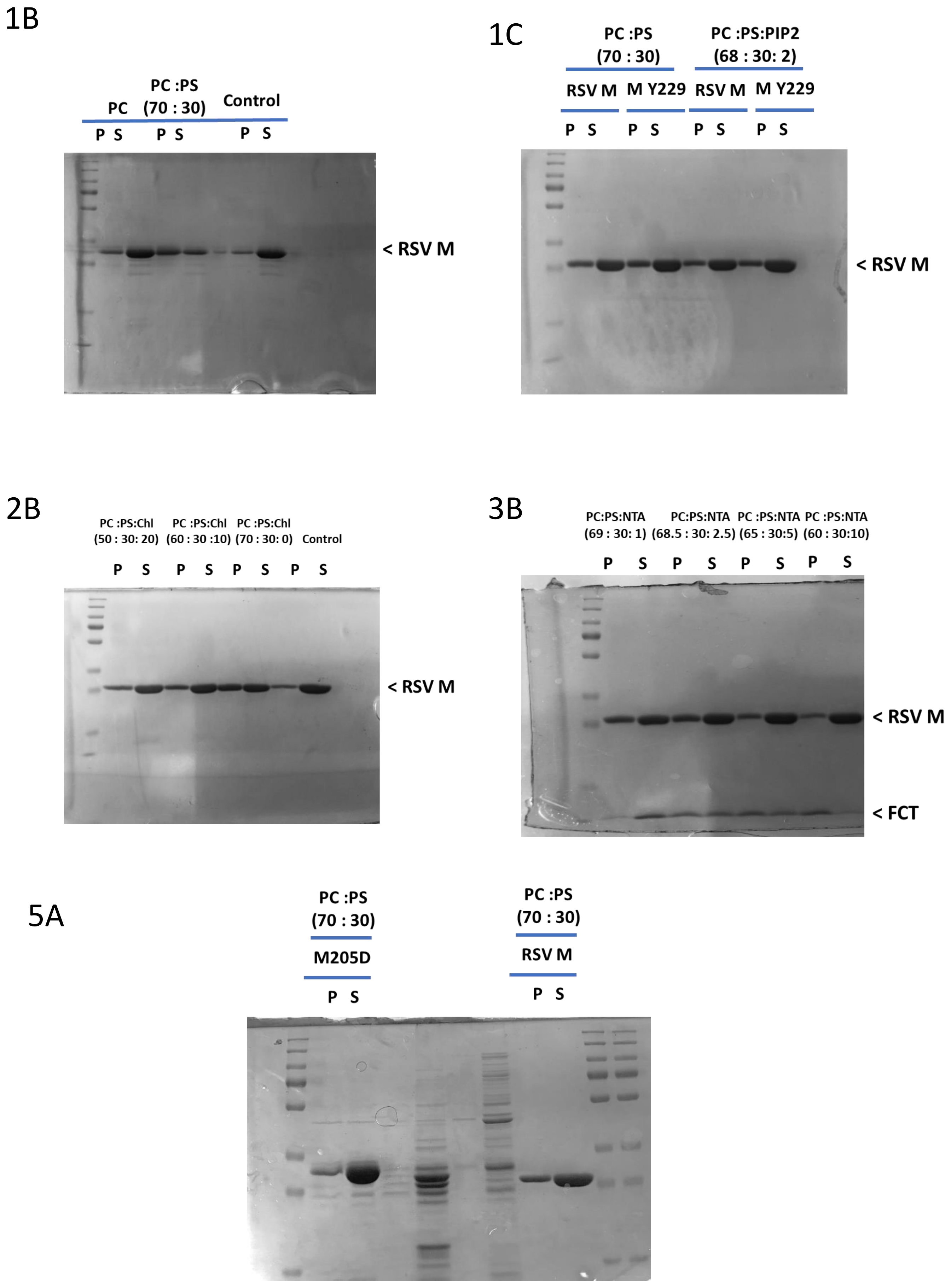


**Supplemental Figure 2**: Full Coomassie gels for 1B, 1C, 2B, 3B, and 5A.
